# Attentional selection drives information convergence within macaque lateral prefrontal cortex

**DOI:** 10.64898/2026.05.05.722905

**Authors:** Daniel J. Mitchell, Mikiko Kadohisa, Makoto Kusunoki, Cheshta Bhatia, Mark J. Buckley, John Duncan

**Author notes:** For the purpose of open access, the author has applied a Creative Commons Attribution (CCBY) license to any Author Accepted Manuscript version arising from this submission.

## Abstract

Attention is a coherent state, in which multiple brain regions converge to represent selected features of a focal object or event. Lateral prefrontal cortex (LPFC), with its flexible coding of multiple task features and their conjunctions, is widely believed to play a key role in this process. While much research has investigated biased competition between features within a brain region, and cofluctuations of activity between regions, less is known about the representational dynamics through which a coherent neural state emerges. Here, we examine directed mutual information concerning multiple feature-specific population codes, between dorsal and ventral LPFC, across three phases of an attentional selection task. We find bidirectional convergence of information regarding multiple task features, but specifically following the period of selection from the visual display. The results show that neural processes driving inter-region coherence are especially salient during a period of cued object selection, despite comparable local information representation during other task phases.

**Significance statement:** In the primate brain, lateral prefrontal cortex (LPFC) is thought to play a key role in selective attention, which is fundamental to goal-directed behaviour, and implies inter-region convergence towards a coherent representational state. Combining neural recordings, multivariate decoding, and analysis of information dynamics, we find local representations of multiple task features during multiple task phases, in separate regions of LPFC, with strong inter-region representational coherence emerging when attention could be directed to a chosen target. Results show that information convergence between these regions is bidirectional, sustained, and reflects multiple features of a chosen object, but, unlike local information representation, is highly specific to the choice phase of the task.

## Introduction

Attention is a coherent state, in which distributed brain regions encode aspects of an attended event or operation. In the primate brain, for example, multiple regions respond to visual objects, each in part coding their different features and implications for behaviour [1–3]. Data from both human [4–6] and monkey [7–13] show that, compared to an unattended object, attended objects produce enhanced neural responses in many brain areas, representing attended and sometimes unattended features of the target object. Visual attention is thus a coherent state in which multiple brain regions converge to process features of the same selected object [14]. More broadly, it is widely proposed that “cognitive control” regions of frontal cortex bias processing across distributed brain regions, assembling the structure of a current cognitive operation [15–17]. A developing attentional focus may also bring coherence within the frontal lobe. In a search task, for example, Kadohisa *et al.* [10] showed that, in the early response to a visual display, neurons in each hemisphere responded largely to the contralateral object, but before the animal’s response, the two hemispheres converged to represent the same, selected target [see also 18, 19].

Such coherence can be achieved in relatively simple connectionist models, based on spreading activation between feature and conjunction units [20–22]. In the primate brain, flexible conjunctive representations are commonly observed within domain-general regions of the frontal and parietal lobes, which exhibit nonlinear mixed-selectivity of individual neurons [23] and adaptive population coding of diverse task features [24–27].

To assess directed communication between regions, popular measures include Granger causality and directed mutual information [28, 29]. These measures quantify how events in one region Y are predicted by past events in another region X, over and above prediction from the past of Y itself. Significant prediction can be interpreted as transfer of information from X to Y, though, alternatively, both X and Y could be reflecting inputs from a third region Z. In the second case, past states of both X and Y could be noisy estimates of the input from Z, so that prediction of the current state in either region is better when considering both past states. In either case, the appearance of significant prediction is consistent with developing coherence, as some influence drives the two regions towards a common state. Typically, such measures are applied to fluctuations in brain activation, demonstrating directed functional connectivity [e.g. 30, 31], but they can also be applied to measures of feature representations, to reveal directed information about specific representational content [32–37].

In a recent study [38], we recorded activity across the ventrolateral (vLPFC) and dorsolateral (dLPFC) prefrontal cortex, as monkeys attended to one object in a visual display. In both regions, in the period following display onset, there was a developing population code for the location of the attended object. Using directed mutual information, we found prediction of future codes in each direction: the strength of the vLPFC code predicted the future of dLPFC, and vice versa, though with a tendency for prediction to be stronger and earlier in the direction vLPFC to dLPFC. Here, we extend these preliminary analyses. In addition to representation of target location, we examine coding of two other features of the task – the identity of the target object, and the identity of the cue that instructed which object to select. We also compare results for the array period, requiring selection of the chosen object, to the preceding cue period and the subsequent post-response period. Strikingly, we find bidirectional prediction for multiple task features, but specifically following the period of selection from the visual display. The results show that neural processes driving inter-region coherence of selected information are especially salient during a period of cued object selection.

## Results

Figure 1 shows the attentional selection task, electrode locations and behavioural performance. In brief: Prior to recordings, animals had learned two sets of four objects, associated with yellow and green cues (Figure 1A). In each session, animals worked through a series of problems based on these object sets. In each problem, one object from each set was the target, selection of this target (by saccade to its location) bringing reward. In a first series of trials (Figure 1B, cycle 1), animals learned the two current targets by trial and error. Subsequently (Figure 1B, cycles 2 - 4), each trial began with a yellow or green cue, indicating which target should be selected. Following the cue period, an array period presented a display of six objects. On receipt of a go signal, the animal made a saccade to one object, receiving reward for selection of the correct target. Data in this paper come just from the cued trials of cycles 2 – 4, i.e. following the period of trial and error learning. [See 39 for analysis of the reinforcement learning process in Cycle 1 trials.] In these cued trials, the overall percentage of correct trials was 79% for one animal (Figure 1D) and 80% for the other (Figure 5B). Accuracy exceeded 60% in cycle 2, increasing to around 90% by cycle 4, suggesting a continuation of learning. The dominant error in all cycles was section of a wrong object associated with the correct cue.

**Figure 1.**
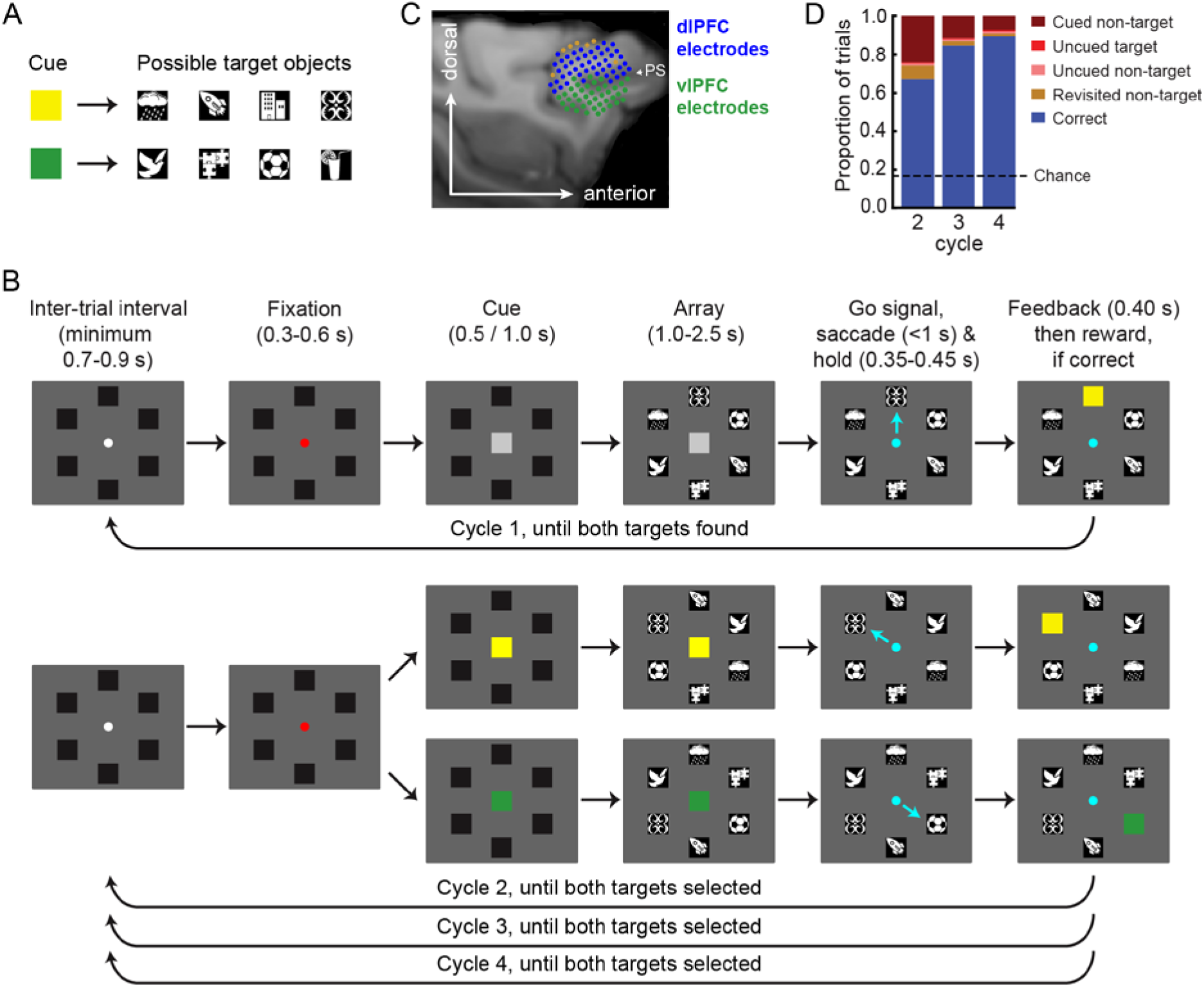
Task, recording sites, and behaviour in animal A. (A) Two possible cues, yellow and green, had previously been associated with four objects each. On each “problem” of the experimental task, the goal was to discover which one of the four possible objects was the rewarded target for each cue, and to select these two rewarded targets across repeated trials. (B) Each row illustrates an example trial and the timings of each phase within a trial. Each problem started with un-cued trials (cycle 1; top row). At this stage, target identities were unknown, and were learned by feedback after each selection. In the feedback period ending each trial, the selected object was replaced by a square. For a correct (target) selection, this square was yellow or green (matching the cue colour associated with this object), accompanied by reward. After incorrect (non-target) selection (or re-selection of a target already rewarded on a previous trial), the square was red and no reward was delivered; after one target had been discovered, accordingly, the animal was to search for the other. Trials repeated, with object locations randomised, until both rewarded targets had been selected. On cycles 2-4, trials were similar, but started with a yellow or green cue, specifying which of the two targets learned in cycle 1 had to be selected on this trial (lower rows). After four cycles of trials, each continuing until each target had been selected once, a new problem started, with a different pair of target objects to be discovered. The examples show feedback squares for correct trials. Cyan arrows indicate example saccades, and were not part of the display. (C) Electrode locations in LPFC of animal A, overlaid on an example sagittal slice of the T1-weighted structural MRI, along with their dominant assignment to dorsal (blue) and ventral (green) regions of interest. Brown electrodes indicate those where no cells were identified in the analysed sessions. Note that region assignments of some electrodes occasionally varied over sessions, as electrodes were advanced in depth. PS, principal sulcus. (D) Behavioural performance of animal A, across non-aborted trials of cycles 2-4.

### Representation of cue identity, target identity, and target location, across the whole trial, in both ventral and dorsal LPFC

We first present results for animal A, from whom the most data were available. Local representation of each task feature by LPFC neural populations is shown in Figure 2. In vLPFC, in line with previous analyses of these data [38], a strong representation of cue identity emerged shortly after appearance of the cue (from about 80 ms), followed by weaker representation of target object identity (from about 200 ms), as cue information began to decline (Figure 2, lower left). Following appearance of the choice array, strong representation of target location emerged (from about 100 ms, Figure 2, lower middle), alongside sustained representations of cue and target identity. Representations of cue, target identity and target location remained significantly above chance until, and beyond, presentation of the post-choice feedback stimulus (Figure 2, lower right).

**Figure 2.**
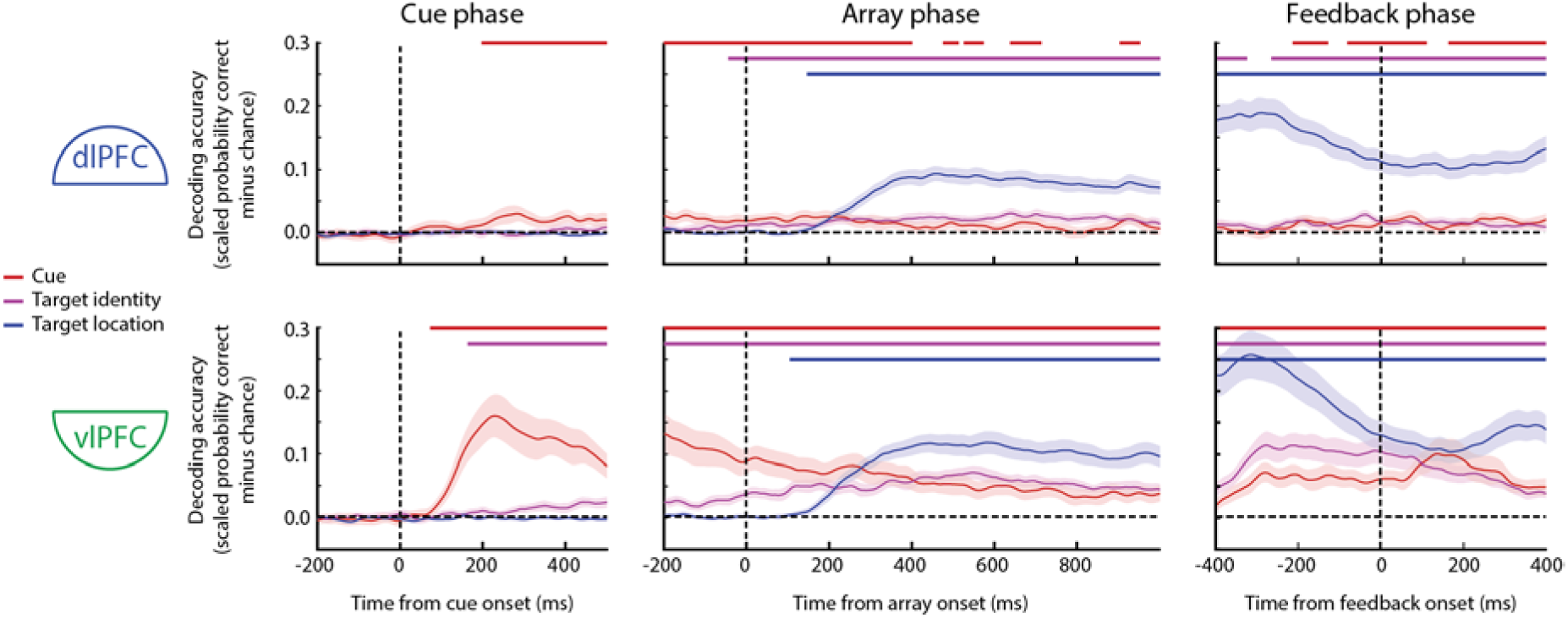
Feature representation within LPFC regions in animal A. Feature decoding accuracies are plotted separately for dLPFC (top) and vLPFC (bottom), and for time points relative to cue onset (left), choice array onset (middle) and feedback onset (right). Accuracy is plotted as the probability of correct classification minus chance, and translucent bands are 95% confidence intervals. Horizontal coloured lines indicate periods of significant representation per feature, corrected for multiple comparisons using TFCE with a maximum statistic permutation test.

In dLPFC, representation of target identity and location followed a similar profile, although identity representation was weaker, and non-significant in the cue phase (Figure 2, upper row). The only substantive difference from the analyses of Kadohisa et al. [38] was that we also found significant, although weak, cue representation in dLPFC, emerging in the cue phase and remaining intermittently significant throughout the array and feedback phases. This difference may stem from the inclusion of additional neurons from the banks of the principal sulcus in the present analyses.

### Bidirectional information convergence across LPFC regions emerges for all features following presentation of the choice array

Previously, using a subset of these data, we have shown that, for target location, significant directed feature information between vLPFC and dLPFC emerged shortly after onset of the choice array [38].

Here, we reiterate this finding and extend it to directed feature information about the identity of the cue and the target object, across all temporal lags (Figure 3A). Directed information concerning all three task features was strong, sustained, and bidirectional, with significant clusters emerging, sharply, from roughly 100 ms after array onset, and persisting beyond the end of the analysis window (1000 ms after array onset). Directed information, especially concerning the target object, was strongest at lags of 50-100 ms, but significant clusters extended up to lags of at least 600 ms. We highlight that transfer of information regarding the cue and regarding the identity of the target did not appear until well after the onset of the choice array, despite significant representation of these features being detectable substantially earlier within both regions.

**Figure 3.**
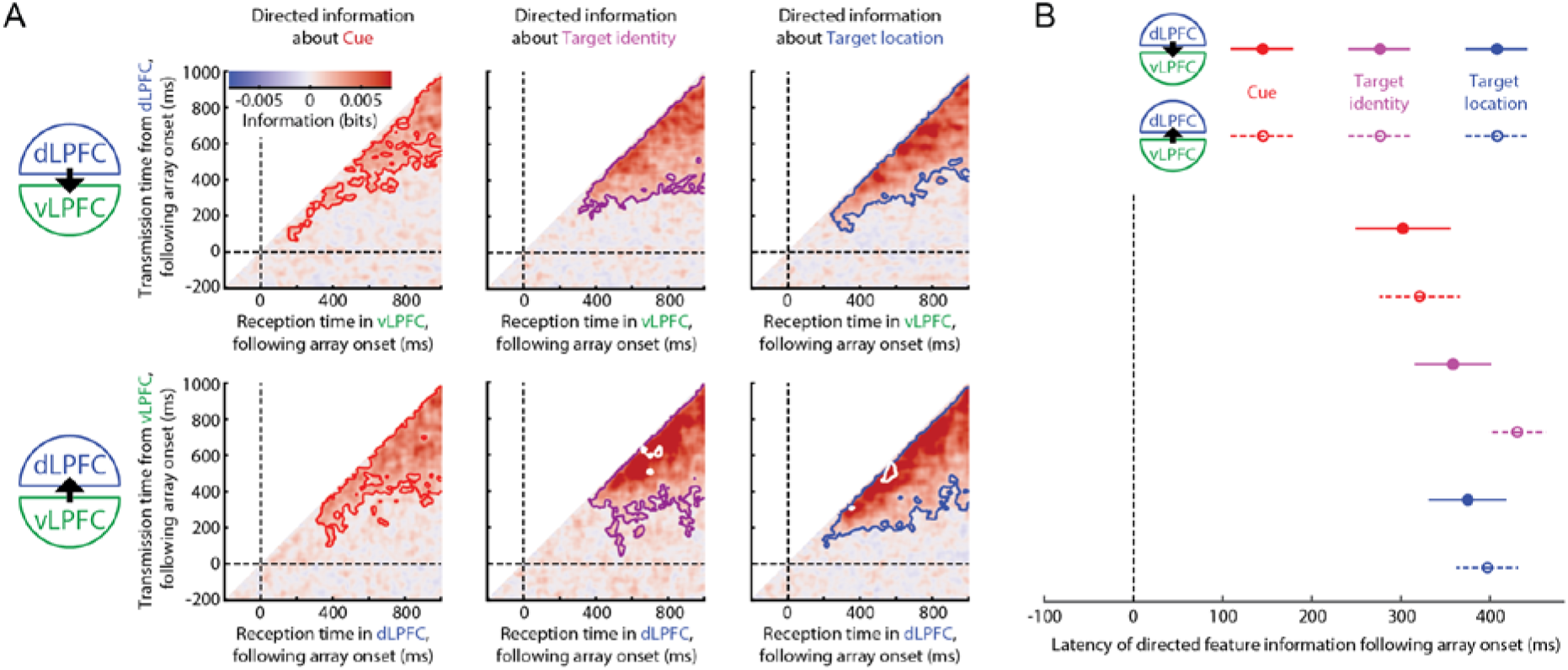
Extensive information convergence across LPFC regions of animal A, following presentation of the choice array. (A) Directed feature information across LPFC, for all pairs of time points relative to array onset, is plotted separately for transmission of information concerning the cue (left), target identity (middle), and target location (right), and separately for transmission from dorsal to ventral (top) and from ventral to dorsal (bottom). Directed information is calculated as conditional mutual information, with bias correction such that chance is zero. Coloured contours indicate periods/lags of significant directed feature information, corrected for multiple comparisons using TFCE with a maximum statistic permutation test. White contours indicate periods/lags when directed feature information is significantly stronger than in the reverse direction, similarly corrected for multiple comparisons. (B) Estimated 25%-area-latency of the onset of directed information about each feature, and in each direction. Error bars are within-session 95% confidence intervals.

There was a general tendency for directed feature information, especially regarding the identity and location of the target object, to be stronger in the ventral-to-dorsal direction; however, this only reached significance at a few time points. Significantly stronger directed feature information in the dorsal-to-ventral direction was not observed.

To compare the latency at which directed feature information began to emerge, for each feature and in each direction, we combined across lags and identified the moment when positive directed feature information exceeded 25% of its cumulative sum across time [40] (Figure 3B). A repeated-measures ANOVA with factors of direction (dorsal-to-ventral, ventral-to-dorsal) and information type (cue, target identity, target location) indicated a main effect of information type (F2,98 = 7.74, p <0.001, BF10 = 21.8), a modest effect of direction (F1,49 = 4.91, p = 0.03, BF10 = 1.64), with earlier transmission in the dorsal-to-ventral direction, but no interaction (F2,98 = 1.23, p = 0.30, BF10 = 0.07). Pairwise t-tests indicated that directed feature information about the cue preceded directed feature information about target identity (t49 = 3.90, p<0.001, BF10 = 87.4) and target location (t49 = 2.76, p=0.008, BF10 = 4.47), with the latter two occurring concurrently (t49 = 0.40, p=0.69, BF10 = 0.17).

### Minimal information convergence is observed in the cue and feedback phases, despite local information representation

Next, we extended the analysis of directed feature information to the other phases of the task. In Figure 4, directed feature information is plotted for the cue phase (left) and for the feedback phase (right). In the cue phase, directed feature information regarding the cue was conspicuously weak, despite representation of cue information in both brain regions being at least as strong as during the array phase. A flicker of directed cue information occurred in the ventral-to-dorsal direction only. The lack of discernible directed feature information concerning target identity is expected, because negligible representation of target identity was measureable within the dLPFC in this period.

**Figure 4.**
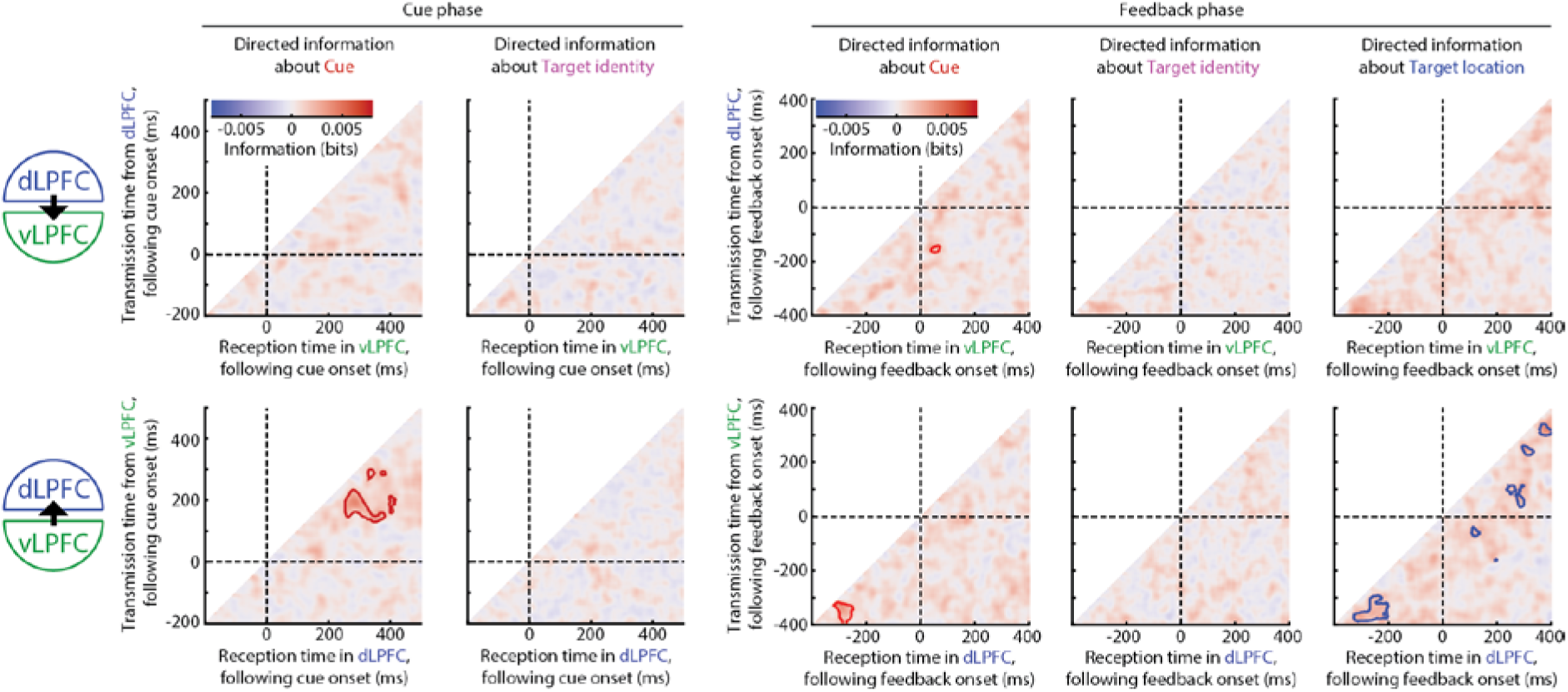
Minimal information convergence across LPFC regions of animal A, following presentation of the cue (left) or the feedback signal (right). Format as in Figure 3.

In the feedback phase, directed feature information regarding the target location was also conspicuously weak, despite representation of location information in both brain regions being substantially stronger than during the array phase. Moments of significant transmission were sparse, and only in the ventral-to-dorsal direction. Directed information regarding the cue was even weaker, and directed feature information regarding target identity was weaker still, again, despite both features being significantly represented in both regions, at a level comparable to the array phase.

### Replication in a second animal

The above analyses were repeated in animal B, and results are plotted in Figure 5. Behavioural performance (Figure 5B) was comparable to animal A. Feature information in both regions largely followed a similar profile to animal A (Figure 5C), but was notably weaker, particularly regarding target identity information, and target location information in the array phase. As for animal A, cue information emerged in both regions shortly after cue onset, remaining significant throughout the trial in the ventral region and intermittently significant in the dorsal region. Unlike animal A, information about target identity did not emerge until the array phase in the ventral region, and until the feedback phase in the dorsal region. As for animal A, information about target location emerged shortly after array onset, and remained significant in both regions throughout the rest of the trial, although at a very low level in the dorsal region during the array phase.

Consistent with animal A, directed feature information about the cue was observed bidirectionally in the array phase, but was not observed in either the cue or feedback phases (Figure 5D), despite moments of cue representation comparable to the array phase in both regions of LPFC. Also consistent with animal A, there was a tendency for directed feature information to be stronger in the ventral-to-dorsal direction, although this was not significant. Unlike animal A, directed feature information concerning the target object (identity and location) was not observed during the array phase (all p<0.05), likely because of minimal representation of these features in the dorsal region. Consistent with animal A, directed feature information about target location was weak (and non-significant) during the feedback phase, despite both regions containing substantial information about target location, at a level much higher than for cue information in the array phase.

### Associations of local and directed feature information with continued learning and with behavioural accuracy

Finally, returning to animal A, we assessed how local feature representation and directed feature information varied with behavioural performance, and across cycles of each problem as learning of the current targets was consolidated (Figure 6). Trials were separated into those with an incorrect choice (“error,” excluded from previous analyses), correct trials following an unrewarded trial (“recovery”) and correct trials following a correct trial (“streak”). On error trials, we continued to decode the correct (unchosen) target. Local and directed information were averaged within time windows of interest, and directed information was averaged across lags.

**Figure 5.**
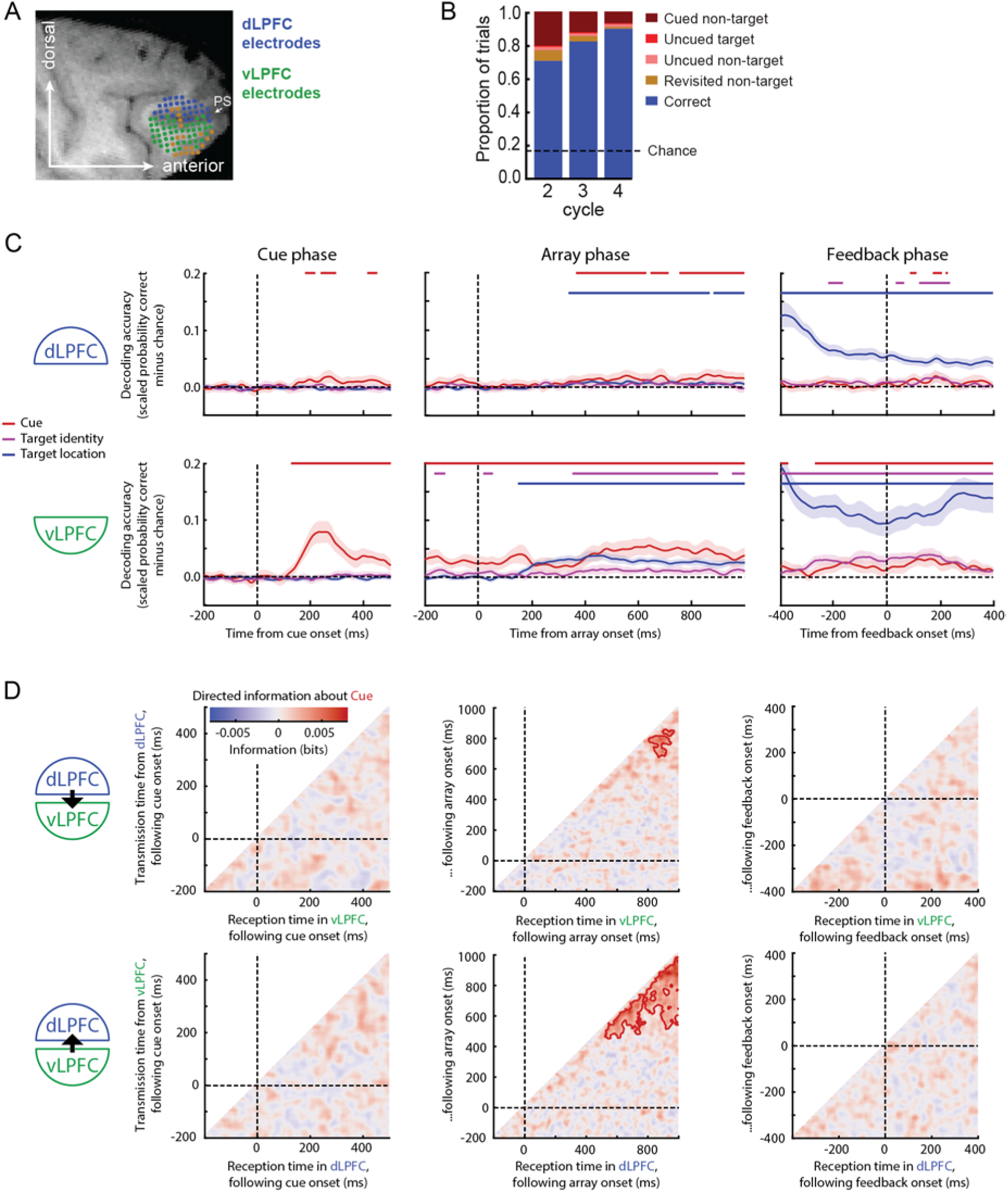
Replication in a second animal, (A) Electrode locations in LPFC, overlaid on an example sagittal slice of the T1-weighted structural MRI, along with their dominant assignment to dorsal (blue) and ventral (green) anatomical regions of interest. Brown electrodes indicate those where no cells were identified in the analysed sessions. Because insufficient cells were recorded on dLPFC electrodes, cells were reassigned per session into more dorsal and more ventral halves. PS, principal sulcus. (B) Behavioural performance across non-aborted trials of cycles 2-4. Chance performance would be 16.7% correct. (C) Feature representation within LPFC regions of animal B. Format as in Figure 2. (D) Convergence of cue information across LPFC regions of animal B, following presentation of the cue (left), choice array (middle) or the feedback signal (right). Format as in Figure 3.

**Figure 6.**
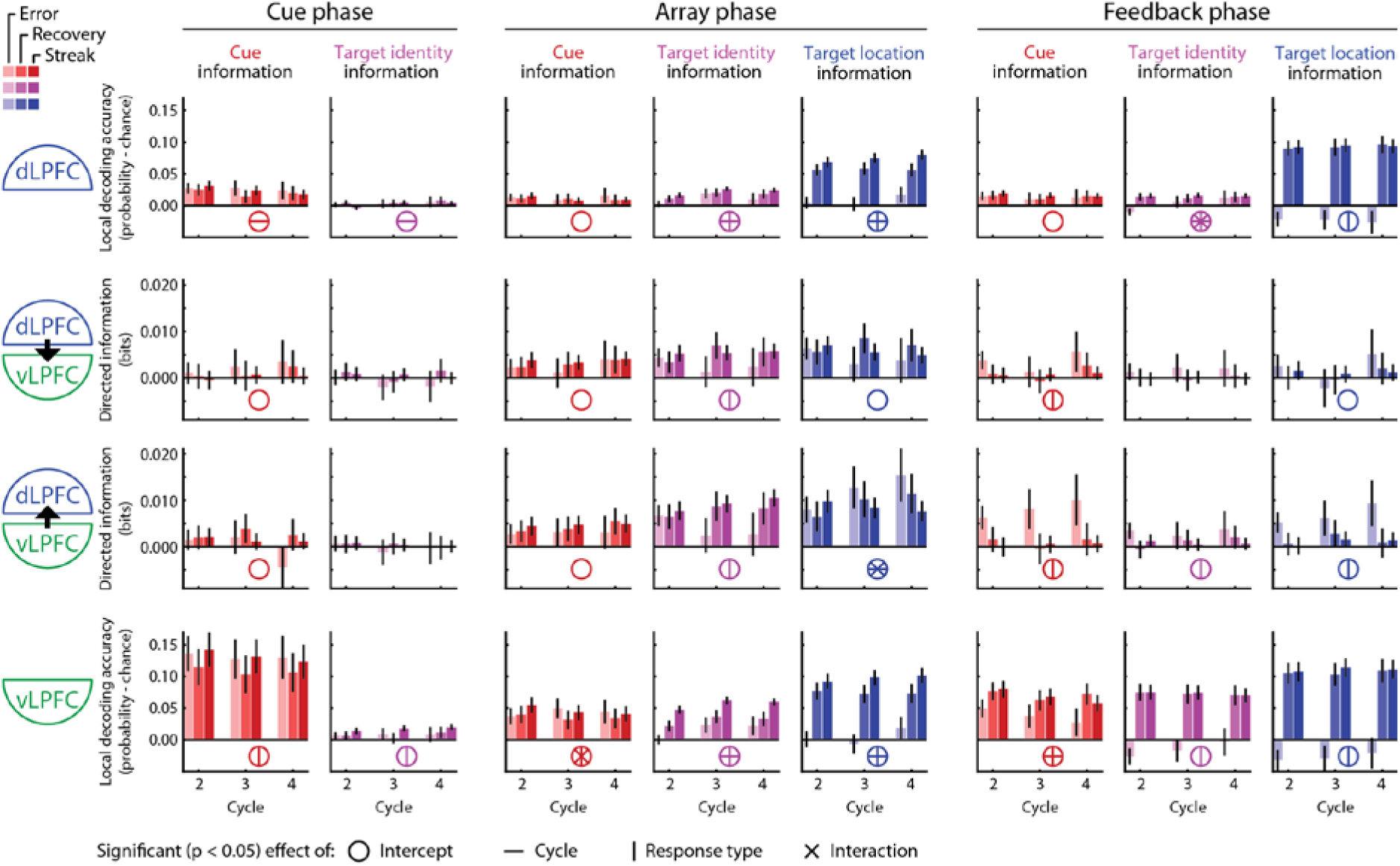
Effects of cycle and behavioural accuracy, in animal A. Local representation of task features (top and bottom rows) and directed information regarding task features (middle rows), broken down by cycle (x axes) and behavioural response (saturation: light = incorrect choice; medium = correct following error; dark = correct following correct). Error bars are 95% confidence intervals.

During the cue phase, vLPFC representations of cue and target identity differed by response but not cycle, with cue representation slightly weaker on trials following errors, and target representation strongest during streaks of correct trials. dLPFC representations of cue and target identity differed by cycle but not response, with cue information slightly weakening while target information slightly strengthened across cycles. Directed information in either direction was weak, regardless of cycle or response.

Following the choice array, local cue information appeared constant across cycle and response types in dLPFC, and modestly depended on their interaction in vLPFC, but without a clear pattern. Local target information (identity and location) in both regions showed independent effects of cycle and response, increasing across cycles, and increasing from errors, to recovery, to correct streaks. Directed information regarding the cue did not significantly vary by cycle or response. Directed information regarding target identity tended to be weaker on error trials. Directed information regarding target location did not vary with cycle or response in the dorsal-to-ventral direction; in the ventral-to-dorsal direction, an interaction suggested stronger cross-region convergence on error trials, but only in later cycles.

At feedback, cue information in dlPFC appeared insensitive to cycle and response, but weakened across cycles in vLPFC, where it was also lowest on error trials. Target representation (identity and location) was strong on correct trials, and generally negative on error trials, consistent with a neural response pattern that differed from the correct target pattern learnt by the classifier. Directed information regarding all features was weak on correct trials, but, interestingly, increased on error trials, perhaps because error feedback triggered a reallocation of attention.

## Discussion

Selective attention is a coherent brain state, in which multiple brain regions converge to encode aspects of the attended event or cognitive operation. Standard measures of directed coherence include Granger causality and directed mutual information, reflecting how the state of region Y is influenced by preceding states of another region X. Here we apply this approach to representational content, to show that, as a target is selected in a multi-object array, directed feature information representing cue and target properties emerges between dorsal and ventral regions of the lateral prefrontal cortex. This inter-region information coherence is sustained throughout the period of selection following presentation of the visual choice array, but, strikingly, appears largely specific to this phase of the task, despite local information representation during other task phases. An exception is increased information convergence also following error feedback. Plausibly, this could also reflect reallocation of attention away from the incorrectly chosen item.

Whenever we observed an asymmetry in the strength of directed feature information, it was stronger in the ventral to dorsal direction than the reverse. From the present data, it is unclear whether this might be because vLPFC has more of a broadcasting role in this task, or simply reflects the fact that measureable representations were generally stronger within vLPFC. In other situations, dominant information transmission in the other direction can be observed [e.g. 34, who examined similar LPFC regions during reinforcement learning]. The dominant direction of information transfer is likely influenced by specific task demands, potentially explaining differences between studies and contexts. More importantly, even in the presence of mild asymmetry, information convergence in the current context appears strongly reciprocal.

The data are not well described by a simple idea of information “transfer” or “flow.” Rather than information being lost from the source region at later times, representational strength was largely sustained in both regions, such that the regions converged on a shared representational state. The observations that directed feature information was sustained and concurrent in both directions lend further weight to the aptness of describing this as a process of convergence. In animal A, the concurrent emergence of directed feature information about both the identity and location of the target is suggestive of an object-based mechanism. Because the target’s identity, retrievable from the cue, specified a search template by which its location could later be selected, the concurrent and relatively late communication concerning both identity and location further suggests that information convergence reflects the outcome of selection, rather than the preceding construction of a search template [15].

### Specificity of information integration to the array phase

The comparison of task phases raises an obvious question: On correct trials, why was information integration observed almost exclusively during the array phase, despite cue information, in both regions, being at least as strong in the cue phase, and location information being at least as strong in the feedback phase? The array phase differs from the cue and feedback phases in comprising a choice between multiple objects, requiring peripheral selection, matching to a search template, and action preparation. In contrast, the cue and feedback phases require attention to a single, binary, foveal cue. Any of these features, or others, could plausibly influence information integration, and we highlight two interesting possibilities.

Firstly, in a dissection of neural components of attention, Wen et al. [41] found that the representation of target identity emerged faster when three objects competed for attention, compared to a single object presented alone. It may be that the presence of multiple competing inputs drives both stronger integration and faster convergence of selective representations.

Alternatively, information convergence across brain regions may be especially important in the context of action selection [42, 43], potentially consistent with recent rodent studies reporting broad propagation of decision-related and motor signals across the brain [44–46]. In the cue phase of the present task, an object representation can be primed, but a physical action cannot yet be planned. Processing the feedback signal involves various cognitive operations, from error monitoring to strengthening associations between cue, object and reward, but, again, without mapping directly to a physical action choice. Greater object-based information convergence in action-oriented cognition could plausibly stem from the physical constraint in fixating one object at a time, consistent with tight coupling between eye movements and selective attention [47]. A dependence on response selection also fits with various findings from the human neuroimaging literature. For example, in frontoparietal regions associated with the LPFC region examined here [48, 49] item features are more strongly represented when priming a specific action rather than simply memorised [50], sensitivity to response competition is greater than to perceptual competition [51], and stimulus-response mappings are represented more strongly than the individual stimuli and responses themselves [52].

One can imagine future experimental manipulations to isolate the necessary and sufficient conditions for information convergence across LPFC. Regardless of the precise boundary conditions, specificity of representational convergence to the choice phase of the task resonates with a recent suggestion that inter-areal communication is more task-state dependent than within-area information processing [30]. Examining cofluctuations of activity, rather than representations of specific attended features, the authors reported that transfer entropy across a macaque frontoparietal grasping network was specifically enhanced during the movement phase of the task, whereas active information storage within each area showed less task-phase dependence. Interestingly, temporary optogenetic silencing of a posterior LPFC region disrupts behavioural performance of macaques when applied during the selection phase, but not during the cue phase of a task [53]. This echoes the current finding of specificity of integration to the array phase, further suggesting a causal role of LPFC representation, and criticality of the selection phase, in supporting goal-directed attention.

### Limitations and generalisability of the current results

Although directed feature information can be considered statistically causal in the sense that information in the “source” region precedes and helps to predict information in the “target” region, it would be premature to conclude direct communication, although this is likely part of the story. It is also possible that directed feature information reflects separate representations of common information from a third input, in the presence of temporal autocorrelation and independent noise. Regardless of this uncertainty, it is clear that processing an object array in this task leads to substantial convergence of selective representations across dorsal and ventral regions of LPFC.

We are limited here to data from two animals, and it would be valuable to confirm that the present findings generalise to other animals and to the human brain. We have also focused on two regions within the lateral prefrontal surface, and object features that are relevant to the task; however, we might expect similar information convergence across much of the brain, including more-domain-specific regions. For example, information regarding the attended hemifield is transmitted from LPFC to area V4 [36]. On the other hand, information integration could be especially strong amongst frontoparietal association regions [54], in part due to their properties of adaptively representing arbitrary task-relevant attributes [24, 25, 27] and conjunctions [23], as well as widespread cortical connectivity [55]. For example, selective attentional convergence might extend beyond the LPFC region considered here to parietal regions, to which LPFC has been shown to transmit attended information [35]. It is also unknown how far information convergence might extend to unattended features of an attended object, when these are represented [56, 57]. Further experiments would be required to determine to what extent the current findings generalise to other brain regions, other object features, and different task settings.

### Conclusion

One can view target decoding within any brain area as a measure of the resolution of attentional competition, and directed feature information as measuring the degree to which this competition is integrated across brain areas. Under this view, the current results suggest that integration, at least across the macaque LPFC, may not be an inevitable consequence of co-representation, but is triggered when one of several objects is selected for action. In this situation, information convergence between ventral and dorsal LPFC is bidirectional, sustained, and can reflect multiple features of the chosen object.

## Methods

### Subjects

Two male rhesus macaques were tested (*Macaca mulatta*, 14 Kg each, aged 7-9 years at test). Experiments were performed in accordance with the UK Animals (Scientific Procedures) Act, 1986, the guidelines of the European Community for the care and use of laboratory animals (EUVD, European Union directive 86/609/EEC), and with ethical approval from Oxford University’s Animal Care and Ethical Review committee. All procedures were licensed by a Home Office Project License.

### Task

The task, illustrated in Figure 1, has been described in detail previously [38]. Prior to the experiment, animals had been trained to associate two possible cues, yellow and green, each with four object images (Figure 1A). Each experimental session was divided into “problems”, each problem having two “target” objects, one from each colour category. On each trial, the animal was shown a display of six objects, and on receipt of a go signal, made a saccade to one of these (“object selection”). Across a first series of trials (Figure 1B, cycle 1), feedback at the end of each trial allowed the animal to discover the two current targets. In subsequent cycles (Figure 1B, cycles 2-4), animals were cued to re-select these same targets for further rewards. Object locations were randomised on each trial, requiring selection by object rather than by location. The task was thus to learn two new targets in cycle 1 of each problem, one from each colour category, then to hold these in working memory for subsequent selection in cycles 2-4.

Each trial (Figure 1B) was preceded by an inter-trial interval, with a central white fixation point surrounded by six black squares (6 x 6 degrees of visual angle, dva), equally spaced around an imaginary circle of radius 14 dva. The minimum duration of this inter-trial interval was 0.7 to 0.9 s; once this delay had passed, the task waited until the eye was centrally fixated (within a window of 2.6 x 2.6 dva for animal A; 3 x 3 dva for animal B), whereupon the fixation point turned red for a further 0.3 to 0.6 s to indicate trial onset. A cue stimulus (also a 6 x 6 dva square) then replaced the fixation point (“cue phase”). After a period of either 0.5 or 1 s (fixed per session), each black square was replaced by a choice object (“array phase”), three objects associated with each cue (one target and two non-targets, omitting the target object from the preceding problem). The animal was to maintain fixation on the central cue for a further 1.0 to 2.5 s, until it was replaced by a cyan fixation point (“go phase”). The animal then had up to 1 s to make a saccade to one of the objects, and to fixate it for 0.35 to 0.45 s. The chosen object was then replaced with a 6 x 6 dva square (“feedback phase”). The animal was to hold fixation for a further 0.4 s, at which point the feedback and object stimuli were removed; on rewarded trials, a drop of soft food was delivered 0.05 to 0.15 s later. Within a cycle, the inter-trial display then followed directly. To mark the end of a cycle, a grey fixation point was presented alone for 3.2 to 3.5 s; to mark the end of a problem, the screen blanked for 3.3 to 3.6 s.

In cycle 1, the cue was light grey (Figure 1B, upper). The animal selected objects by trial and error. The first selection of either of the two current targets led to positive feedback (yellow or green square, corresponding to the colour category of the selected target) and reward. Any other selection led to negative feedback (red square) and no reward. In this cycle, repeated selection of an already- rewarded target also led to a red square and no reward; after one target had been discovered, accordingly, the animal was to search for the other. Trials repeated until both rewarded targets had been selected, in either order. In each of cycles 2-4, the cue was either yellow or green, indicating which of the two learned targets should be selected. The cue colour was randomised until the first target was selected, after which only the other cue colour was presented. Selection of the cued target led to a feedback square of the corresponding colour and reward; any other selection led to red square and no reward. Optimally, therefore, each of cycles 2-4 consisted only of two trials, one for each target.

If fixation was broken prior to the go signal, or prior to the feedback stimulus, the trial was aborted immediately without reward, and repeated with the same cue and object positions. Aborted and subsequently repeated trials were excluded from analyses, along with incorrect trials.

Stimuli were presented on a 17.5-inch LED screen, controlled using REX data acquisition software (version 8; National Institutes of Health). Eye position was recorded using a 120 Hz, infrared eye-tracking system (Applied Science Laboratories).

### Neural recordings

As described previously [38], each subject was implanted with a titanium head holder and semi- chromic, microdrive recording chambers, with 1.5 mm inter-electrode spacing (form-fitting chamber system, Gray Matter Research). A craniotomy under each chamber enabled physiological recordings. Surgery was performed in aseptic conditions, under general anaesthesia. 96-channel, frontal chambers were positioned over the lateral prefrontal cortex of the right hemisphere (target position: x = 14.7 mm, y = 34.9 mm for monkey A; x =19.1 mm, y = 36.4 mm for monkey B), anterior to the arcuate sulcus, and spanning the principal sulcus (Figure 1C). A chamber targeting temporal regions was also placed over the right parietal cortex, but is not analysed here.

Neural data were recorded, amplified, and filtered (300 Hz to 10 kHz) using Cereus System (Blackrock Microsystems). Cluster separation, to isolate neural activity, was performed with Offline Sorter (versions 3.3.5 & 4.6.2, Plexon Inc.). Between each session, electrodes were independently advanced by a minimum of 62.5 μm, until activity could be identified from new cells.

Data were available from 122 daily sessions for animal A, and 125 for animal B, with an average of 57 completed problems per session.

### Pre-processing of neuronal signals

The first goal of analysis was to obtain time courses of decoded stimulus information, per trial, from the population response in each of dLPFC and vLPFC (see Figure 1C for electrode locations in animal A, and Supplementary Figure 1 for a schematic overview of the analysis approach). Recorded neurons were localised based on post-mortem histology and MRI. For the primary analysis in animal A, data were analysed from all sessions (n=50) in which at least ten neurons were simultaneously recorded from both dLPFC (defined as area 46d and lateral parts of areas 8 and 9 adjacent to this) and vLPFC (defined as areas 46v, 45a, 12l, and 12r). Note that we included neurons from the banks of the principal sulcus, which had been analysed separately by Kadohisa et al. [38]. Including these neurons allowed us to include a larger number of sessions that met the arbitrary criterion of ten or more neurons per region. For animal B, only five sessions contained sufficient neurons in both anatomically-defined regions, according to the above criteria. To allow a reasonable number of sessions from animal B, we instead included sessions with at least 16 neurons across the whole LPFC, and divided these neurons into a dorsal set and a ventral set, of approximately equal size, per session. We excluded four sessions with fewer than 150 correct trials at cycles 2-4, leaving 47 analysed sessions from animal B.

All analyses were restricted to correct trials from cycles 2-4 of each problem (Figure 1D), when the rewarded target objects were known, and explicit cues were presented, so that cue and target object identity information would be available upon presentation of the cue. Across analysed sessions, the percentage of correct trials on cycles 2-4 ranged from 65.7% to 91.8% (mean 79.2%) for animal A, and 68.7%-87.0% (mean 79.8%) for animal B. The number of analysed trials per session ranged from 243 to 370 (mean 319.0) for animal A, and 160-336 (mean 237.2) for animal B.

Following the observation in Kadohisa et al. [38] that neuronal firing rates and high-frequency power (HFP) of local field potentials (LFPs) carry partially independent information about trial variables, we extracted both signals. We assume that the HFP reflects the combined activity of multiple neurons in the vicinity of each electrode [58, 59]. All analyses were performed using Matlab (version 2018a; The MathWorks), having loaded the data using the Neural Processing MATLAB Kit (version 5.5.2.0, Blackrock Microsystems).

Per session, LFPs were analysed from any electrode at which neurons had been recorded. using the Fieldtrip toolbox [60], LFPs were first down-sampled from 30 kHz to 5 kHz and then to 1 kHz, and notch filtered at 50 Hz and its harmonics, to suppress power line noise. To identify noisy trials and electrodes, data were temporarily epoched to the time-window of interest (see below), and bad trials per electrode were defined as those whose signal deviated by more than six standard deviations from the mean over time. Electrodes with >20% bad trials were excluded (zero for animal A; one for animal B), and remaining electrodes were re-referenced by subtracting the mean signal across electrodes per region. Remaining bad trials were excluded (animal A: median 0.6 %, range 0-15.9 %; animal B: median 0.9 %, range 0-14.6 %), and instantaneous HFP was extracted by band-pass filtering between 70-250 Hz and taking the square of the Hilbert transform.

To extract the firing rate of directly recorded neurons, binary vectors of action potentials, sampled at 1 kHz, were smoothed with a Gaussian kernel with a full-width-half-maximum of 50 ms. Similar smoothing was applied to the HFP time-series.

Finally, both signals were epoched into time windows around each event of interest (-200 to +600 ms around cue onset; -200 to +1000 ms around array onset; -600 to +600 ms around feedback onset), and further down-sampled to a temporal resolution of 2 ms.

### Local Feature representation

Separately per region (dLPFC, vLPFC), per session, and per time point, a linear discriminant classifier was used to decode each feature of interest (cue identity, target identity, target location), using 5-fold cross-validation across trials, a uniform prior, and a diagonal covariance matrix. Each classifier used firing rates of all recorded neurons, and HFP of selected electrodes as described above, because we have shown previously that these signals provide complementary information to the classifier [38]. For each time point of each trial, the classifier returned the probability that the predicted class matched the true class, i.e. an estimate of information that the neural population carried about the classified feature.

Each cue was associated with a fixed set of four objects, so cue decoding encompasses information about the cue itself and its set of associated objects. Decoding of target object identity was performed within each set, and then averaged, to avoid any contribution from the cue.

To assess decoding accuracy, performance was summarised per session as the mean probability of correct classification minus chance, averaged first across trials per class, and then across classes, and scaled so that a value of 1 would indicate perfect classification. Above-chance decoding was then tested using a one-sample t-test against zero, across sessions per time point. Correction for multiple comparisons was performed using Threshold-Free Cluster Enhancement [TFCE; 61], with default parameters (height exponent=2, extent exponent=0.5). The family-wise error rate (FWER) was controlled by comparing the statistic at each time point to the 95^th^ percentile of a null distribution of the maximal statistic across time, constructed from 1000 permutations with random sign flipping.

### Directed feature information

For each task feature, we estimated cross-region directed information (DI) [29, 62], by calculating the time-lagged conditional mutual information between the single-trial time-series of feature representation estimates described above. Specifically, for a pair of past and future time points, separated by a given lag, we estimated the mutual information between the trial-wise vectors of feature information estimates from the past of the “source” brain region and the future of the “target” brain region, conditional on the past of the target region. DI is also known as transfer entropy [29, 63, 64], and is an information theoretic analogue of Granger causality [29, 65, 66], which, when applied to time-courses of representational strength, can be interpreted as “directed feature information”.

DI was calculated following Gaussian copular normalisation of the input vectors, and with bias-correction of the estimated entropies, as implemented in the Gaussian-Copula Mutual Information Toolbox [29]. The copular transformation provides an efficient lower-bound estimate on the mutual information, and the bias correction ensures that the DI estimates have an expectation of zero under the null hypothesis of no relationship between the regions [see 29, for further details on the benefits of this estimation approach]. We implemented this method of estimating DI (i.e. transfer entropy) within the framework of Integrated Information Decomposition [67, 68], obtaining identical values to those produced by the Gaussian-Copula Mutual Information Toolbox [29].

For each pair of time points, a one-sample, one tailed, t-test across sessions was used to test whether bias-corrected DI estimates were above zero. Significance was corrected for multiple comparisons using TFCE (default parameters; permutation test of the maximum statistic across pairs of time points, against a null distribution from 1000 permutations with random sign flipping). Paired, two-tailed t- tests were used to compare the strength of DI in the two directions, with significance similarly corrected for multiple comparisons using TFCE and 1000 sign permutations.

### Estimating onset latency of directed feature information

To estimate, per session, the latency of emerging DI regarding each feature, we first combined across lags by reflecting each DI matrix about its diagonal, and then averaging down the columns. Then, we used 25%-signed-area latency to estimate the rise-time of the DI distribution [40]. Thus, for each session, onset latency was taken as the moment (per feature and direction) at which the summarized, positive DI first exceeded 25% of its cumulative sum across time.

Onset latencies were compared across features and directions using across-session repeated-measures ANOVA, with Greenhouse-Geisser adjustment for non-sphericity, followed up with paired t-tests. Frequentist statistics were supplemented with default Bayes factors based on the F statistics [69] and t statistics [70].

### Associations of local and directed feature information with continued learning and with behavioural accuracy

In animal A, to assess consolidation of learnt cue-target associations per problem, trials were separated across cycles 2-4. To test associations with behaviour, trials were further separated into those with an incorrect choice (“error,” excluded from previous analyses), correct trials following an unrewarded trial (“recovery”) and correct trials following a correct trial (“streak”). When calculating local feature representation, the training set for each classifier included trials from all categories, and classifier performance was measured separately on held-out test trials from each combination of cycle and behaviour category. DI was then calculated from these condition-specific classification accuracies, per session.

Local and directed feature information were averaged across time windows of interest (and lags, for directed information): 200-500 ms post cue, 400-1000 ms post choice array, and -200 to 200 ms around feedback. Then, for local feature information in each region, and DI in each direction, main effects of cycle, behavioural accuracy, and their interaction were assessed using linear models with session as random effect. Because low numbers of trials for some factor combinations (e.g. errors in cycle 4) increased the variance of the DI estimate [29], observations were weighted by the number of trials.

### Data availability

The analysis code and epoched data (neuronal firing rates, high-frequency power and event files) underlying this article are available on the Open Science Framework (https://osf.io/bfdz9/).

## Acknowledgements

This work was supported by Wellcome Trust grant 101092/Z/13/Z and Medical Research Council Intramural Program MC_UU_00030/7. We are grateful to Andres Canales-Johnson for helpful discussion. The authors declare no competing financial interests.

## Author contributions

J.D., M.J.B. M. Kadohisa, M. Kusunoki, and D.J.M. designed the research. M. Kadohisa and M. Kusunoki collected the data. D.J.M., M. Kadohisa, M. Kusunoki, and C.B. analyzed the data. D.J.M. wrote the paper. D.J.M. and J.D. revised the paper.

## Supplementary material

**Supplementary Table 1.** Numbers of analysed channels and neurons across sessions in each area.

| Animal | Area | Number of channels |  | Number of neurons |  |
| --- | --- | --- | --- | --- | --- |
|  |  | Median | Range | Median | Range |
| A | vLPFC | 17 | 9-31 | 21 | 10-55 |
|  | dLPFC | 16 | 8-33 | 21 | 10-63 |
| B | vLPFC | 8 | 5-23 | 9 | 8-29 |
|  | dLPFC | 9 | 6-24 | 10 | 7-29 |

**Supplementary Figure 1.**
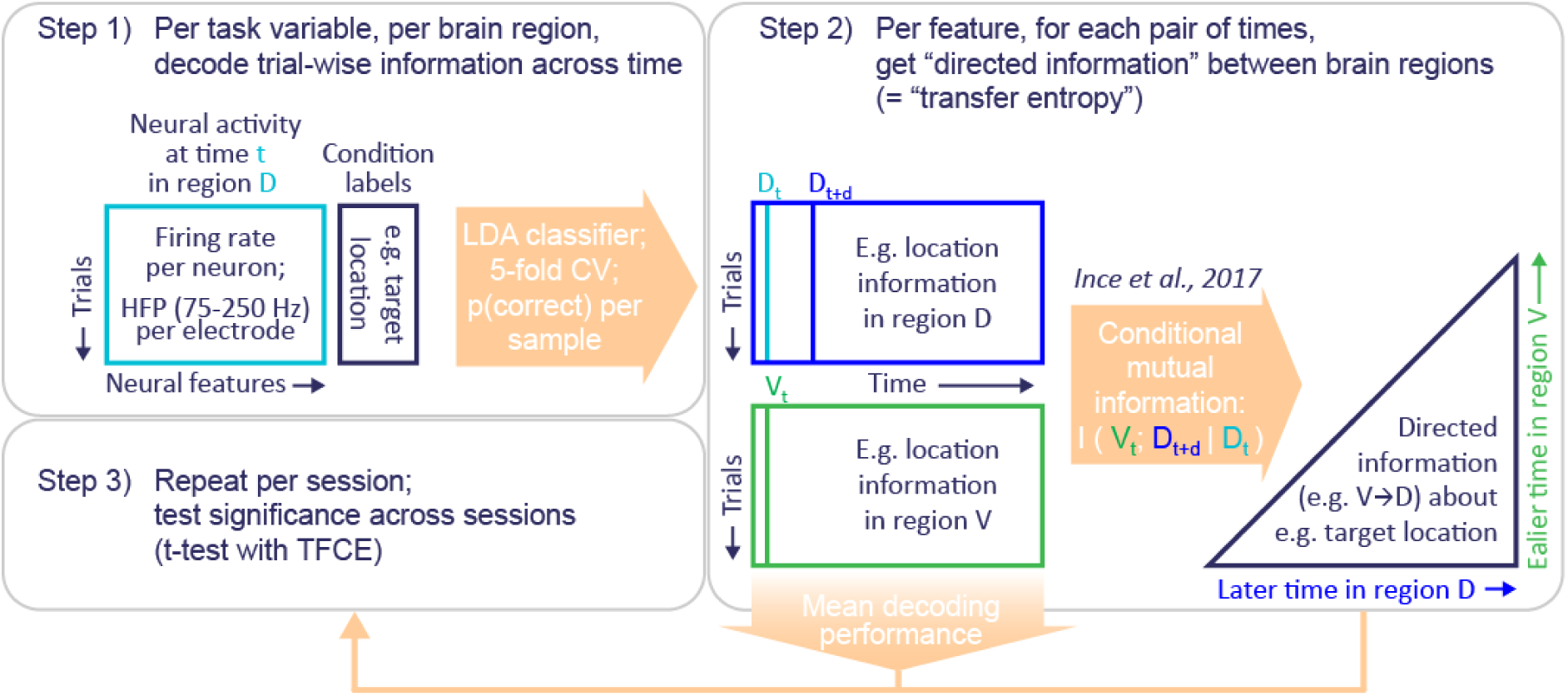
Schematic overview of the analysis approach. HFP = high-frequency power; LDA = linear discriminant analysis; CV = cross-validation; TFCE = threshold-free cluster enhancement.

